# *Pseudomonas aeruginosa* reverse diauxie is an optimized, resource utilization strategy

**DOI:** 10.1101/2020.07.28.224436

**Authors:** S. Lee McGill, Yeni Yung, Kristopher A. Hunt, Michael A. Henson, Luke Hanley, Ross P. Carlson

## Abstract

*Pseudomonas aeruginosa* is a globally-distributed bacterium often found in medical infections. The opportunistic pathogen uses a different, carbon catabolite repression (CCR) strategy than many, model microorganisms. It does not utilize a classic diauxie phenotype, nor does it follow common systems biology assumptions including preferential consumption of glucose with an ‘overflow’ metabolism. Despite these contradictions, *P. aeruginosa* is competitive in many, disparate environments underscoring knowledge gaps in microbial ecology and systems biology. Physiological, omics, and *in silico* analyses were used to quantify the *P. aeruginosa* CCR strategy known as ‘reverse diauxie’. An ecological basis of reverse diauxie was identified using a genome-scale, metabolic model interrogated with *in vitro* omics data. Reverse diauxie preference for lower energy, nonfermentable carbon sources, such as acetate or succinate over glucose, was predicted using a multidimensional strategy which minimized resource investment into central metabolism while completely oxidizing substrates. Application of a common, *in silico* optimization criterion, which maximizes growth rate, did not predict the reverse diauxie phenotypes. This study quantifies *P. aeruginosa* metabolic strategies foundational to its wide distribution and virulence.

## Introduction

*Pseudomonas aeruginosa* is an opportunistic pathogen commonly isolated from diabetic ulcers, burn wounds, and battlefield injuries, as well as from the lungs of patients with cystic fibrosis (CF)[1-3]. Its presence is correlated with high patient morbidity and mortality [4-6]. *P. aeruginosa* is found in ∼ 80% of chronic, diabetic ulcers which cost the US medical system $20-50 billion per year to treat [4-6]. *P. aeruginosa* virulence and persistence mechanisms are enabled by a genome possessing approximately 5,200 core genes, one of the largest of any bacterium [7, 8]. Maintaining the large genome and implementing the myriad of virulence strategies necessitates effective metabolic strategies for nutrient acquisition and allocation. While foundational to its global distribution and virulence, the basis of the *P. aeruginosa* central metabolism is poorly understood [4, 6].

Global regulatory systems select preferred carbon sources from pools of substrates in a process known as carbon catabolite control (CCC) or carbon catabolite repression (CCR) [9]. The best studied examples of CCR are from *Escherichia coli* and *Bacillus subtilis* [9-14]. The metabolic designs of these model organisms, which prefer glucose over other substrates, form the basis of most textbook CCR examples [15]. The CCR strategy represented by *E. coli* and *B. subtilis* is referred to here as ‘classic carbon catabolite repression’ (cCCR) to distinguish it from the broader CCR term. *P. aeruginosa* does not display a cCCR phenotype. Instead, this competitive microorganism, as evidenced by its global distribution which is arguably broader than *E. coli* [16-18], has substrate preferences that are nearly opposite of *E. coli. P. aeruginosa* utilizes a CCR strategy termed ‘reverse diauxie’ or reverse CCR (rCCR) because the hierarchy of preferred carbon sources is nearly reverse that of cCCR preferences [9,33]. *P. aeruginosa* can readily catabolize glucose although it is not a preferred substrate, instead this bacterium preferentially catabolizes less energetic, nonfermentable substrates like succinate. The contrarian hierarchy of preferred carbon sources is proposed to be central to the versatility of *P. aeruginosa*. The ecological basis of rCCR is an open question with few published theories [9, 10, 19-21]. A quantitative understanding of rCCR lags cCCR. This is a critical knowledge gap that contributes to degradation of patient quality of life and costs society tens of billions of dollars a year [5].

Natural environments do not permit unconstrained microbial growth [22]. Instead, life is constrained by the availability of resources such as reduced carbon or nitrogen sources [22]. Phenotypic plasticity can permit microorganisms to acclimate to resource scarcity [23-25]. *In silico* systems biology approaches have investigated resource investments (e.g. carbon, nitrogen) into different metabolic pathways via the enzyme synthesis requirements. The *in silico* methodologies, often referred to as resource allocation analysis or metabolic tradeoff theory, are powerful tools for predicting and interpreting phenotypes and have been applied extensively to cCCR microorganisms *E. coli* and *B. subtilis* [23, 26-33]. For example, *in silico* and *in vitro* studies of *E. coli* quantified acclimation to carbon, nitrogen, or iron limitation along a metabolic tradeoff surface by optimizing the functional return on the limiting nutrient, at the expense of substrates found in excess [25, 34]. This strategy resulted in ‘overflow metabolisms’ with the secretion of byproducts like acetate and lactate; overflow metabolisms are also known as the Warburg or Crabtree effect in eukaryotes [35]. Resource allocation analysis has not been applied to rCCR organisms. Given the large genomic potential and phenotypic plasticity of *P. aeruginosa*, these approaches hold potential for decoding the metabolic organization of this problematic bacterium.

Here, the ecological basis of *P. aeruginosa* rCCR was tested using a combination of physiological studies, exometabolomics, proteomics, and systems biology. The preference for substrates was measured and phenotypic characteristics, like the general lack of an overflow metabolism, were quantified. Proteomics measured a constitutive core metabolism centered on respiration and a dynamic set of enzymatic pathways that catabolized specific substrates, directing intermediates toward the core metabolism. The experimental data was analyzed with a genome-scale, metabolic model of *P. aeruginosa* to identify ecological theories that predicted the observed phenotypes. *P. aeruginosa* did not optimize substrate preference based on standard systems biology assumptions such as the maximization of growth rate, as is commonly applied to cCCR phenotypes. Instead, *P. aeruginosa* metabolism was organized around a multidimensional, resource utilization strategy with constitutive expression of a respiration-based, core metabolism and substrate preferences that were based on minimizing the nutrient investment required to completely oxidize the substrate. Understanding a molecular-level basis of substrate preference, energy metabolism, and cell growth is foundational to controlling virulence mechanisms in *P. aeruginosa*.

## Results

### Growth Physiology and Substrate Preference of rCCR

*Pseudomonas aeruginosa* strain 215 (Pa 215) is a medical isolate from a chronic wound [36, 37]. Pa 215 was grown in chemically-defined, CSP G medium (materials and methods, supplemental material S1). Cultures exhibited two distinct exponential growth phases followed by stationary phase (Fig. 1). A subset of amino acids was consumed preferentially during the first exponential growth phase, which had the highest specific growth rate (Fig.1, Table 1). The second exponential growth phase corresponded with the catabolism of second and third tier amino acids and glucose (Table 2). The cultures did not exhibit an overflow metabolism defined by the secretion of reduced metabolic byproducts like acetate, as is typical of microorganisms expressing cCCR phenotypes [21]. Trace amounts of gluconate were secreted during glucose metabolism but were quickly depleted (supplemental material S2).

**Table 1.**
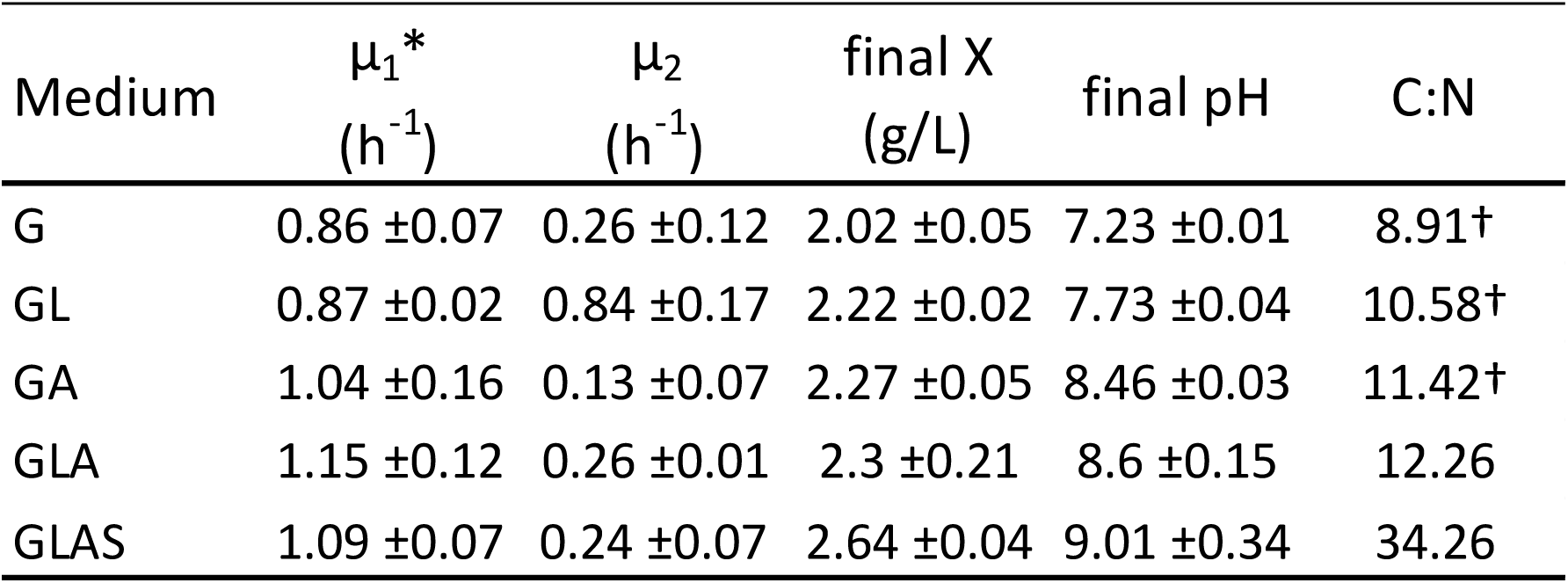
Physiological parameters for *P. aeruginosa* 215 cultures grown on five different, chemically-defined media supplemented with different carbon sources. G = glucose, GL = glucose and lactate, GA = glucose and acetate, GLA = glucose, lactate, and acetate, GLAS = glucose, lactate, acetate, and succinate. Cultures had two exponential growth phases with different specific growth rates (μ_1_, μ_2_). X = dried biomass concentration (g/L). Initial culture pH was 7.0. C:N = ratio of total moles of carbon to total moles of nitrogen in the medium. See text for more details. *The specific growth rate during the first exponential growth phase was not significantly different between condition G and the four other conditions (p-values all > 0.05 using a paired two tailed t-test). ^†^Carbon limited medium. All specific growth rates were calculated from three biological replicates during exponential growth. Final pH values and final biomass values are averages of three biological replicates.

| Medium | $\mu_1^*$<br>(h<sup>-1</sup>) | $\mu_2$<br>(h<sup>-1</sup>) | final X<br>(g/L) | final pH | C:N |
| --- | --- | --- | --- | --- | --- |
| G | 0.86 ±0.07 | 0.26 ±0.12 | 2.02 ±0.05 | 7.23 ±0.01 | 8.91† |
| GL | 0.87 ±0.02 | 0.84 ±0.17 | 2.22 ±0.02 | 7.73 ±0.04 | 10.58† |
| GA | 1.04 ±0.16 | 0.13 ±0.07 | 2.27 ±0.05 | 8.46 ±0.03 | 11.42† |
| GLA | 1.15 ±0.12 | 0.26 ±0.01 | 2.3 ±0.21 | 8.6 ±0.15 | 12.26 |
| GLAS | 1.09 ±0.07 | 0.24 ±0.07 | 2.64 ±0.04 | 9.01 ±0.34 | 34.26 |

**Table 2.**
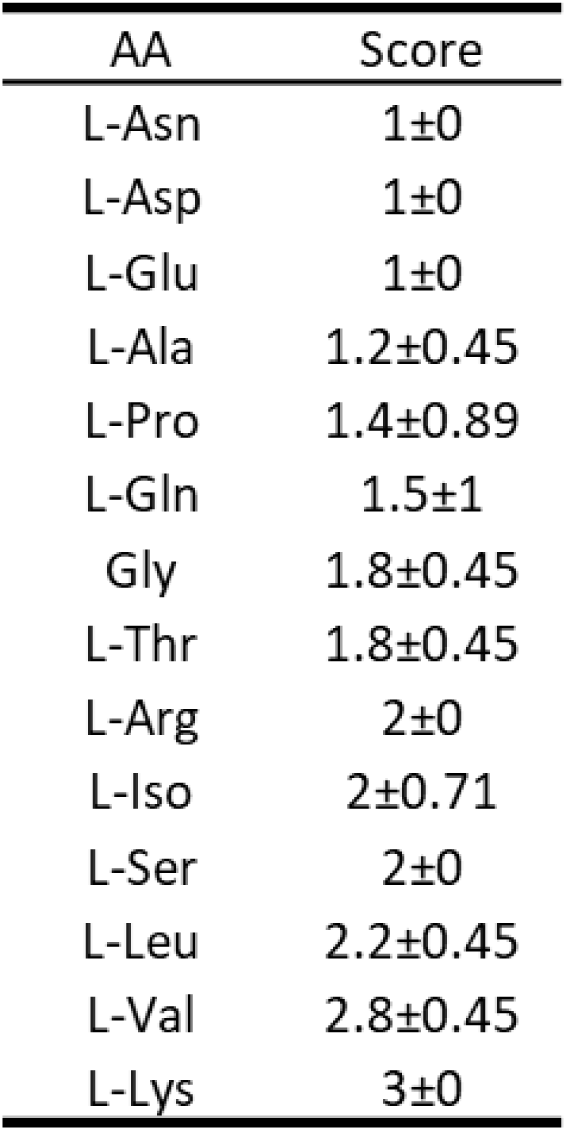
Amino acid preference for *P. aeruginosa* 215 cultures grown on five different, chemically-defined media supplemented with various additional carbon sources. Amino acids were binned into three categories (1,2,3) based on the time of exhaustion averaged between three biological replicates for each of five medium conditions, n=15.

| AA | Score |
| --- | --- |
| L-Asn | 1±0 |
| L-Asp | 1±0 |
| L-Glu | 1±0 |
| L-Ala | 1.2±0.45 |
| L-Pro | 1.4±0.89 |
| L-Gln | 1.5±1 |
| Gly | 1.8±0.45 |
| L-Thr | 1.8±0.45 |
| L-Arg | 2±0 |
| L-Iso | 2±0.71 |
| L-Ser | 2±0 |
| L-Leu | 2.2±0.45 |
| L-Val | 2.8±0.45 |
| L-Lys | 3±0 |

**Figure 1.**
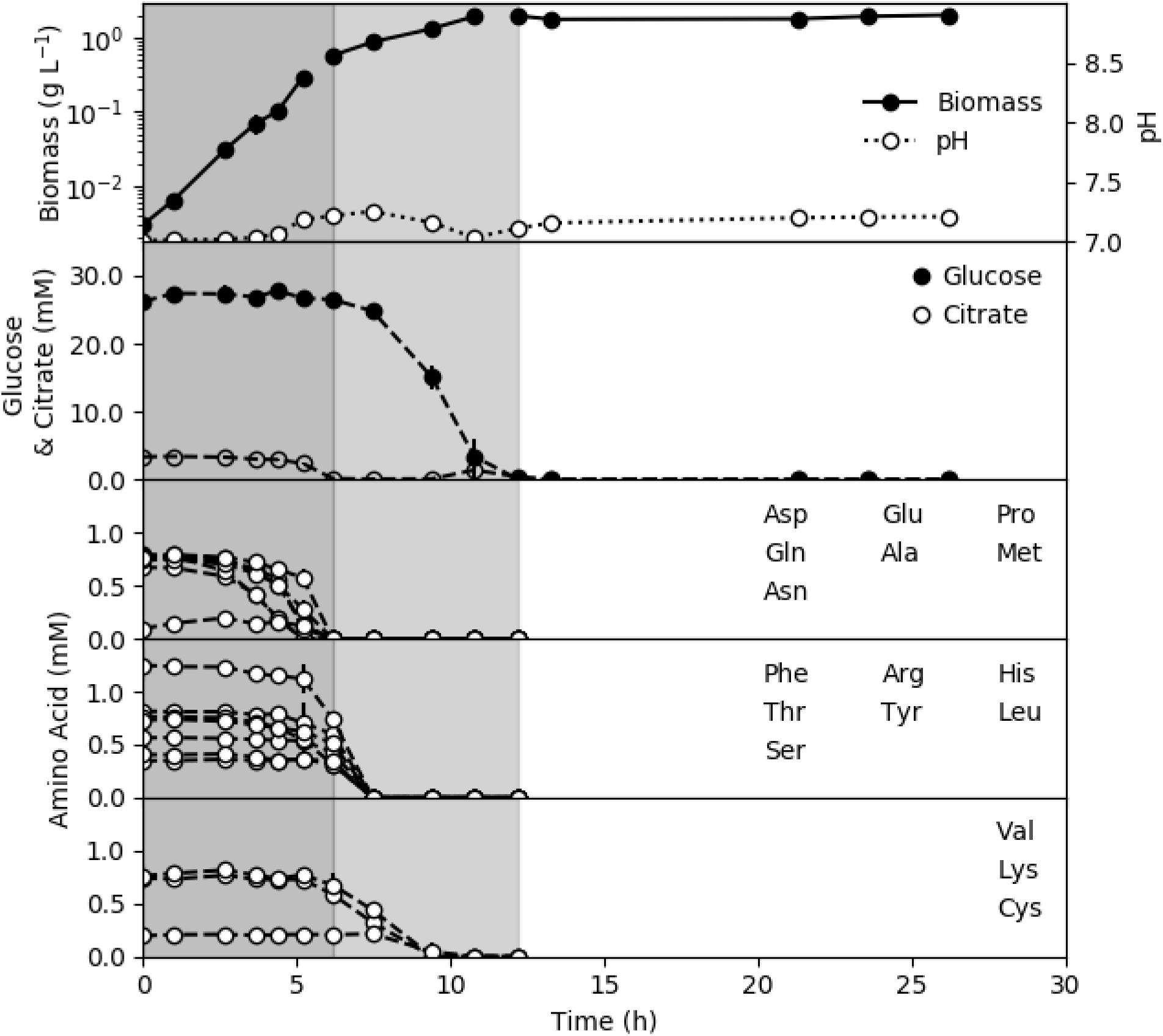
*P. aeruginosa 215* was grown in chemically-defined medium (CSP G) in batch culture. Cultures demonstrated two exponential growth phases highlighted with different background shading. Amino acids are binned into three categories based on their time of exhaustion. Top tier amino acids were consumed during the first exponential growth phase while lower tier amino acids and glucose were consumed during the second exponential growth phase. All values are averages of three biological replicates, and metabolite values are also averaged from two technical replicates. Data can be found in the supplemental material.

Substrate preferences for Pa 215 were quantified using five different formulations of CSP G medium supplemented with permutations of additional carbon sources: lactate (L), acetate (A), and succinate (S) (Fig. 2). Pa 215 grown on CSP GL medium, preferentially consumed the top tier amino acids, represented by aspartate in Fig. 2, followed by lower tier amino acids and lactate before finally catabolizing glucose. Glucose catabolism was not observed while lactate was present. Pa 215 grown on CSP GA consumed the top tier amino acids followed by lower tier amino acids and acetate and finally glucose after the acetate was exhausted. Pa 215 grown on CSP GLA preferentially consumed top tier amino acids, then lower tier amino acids and lactate followed by acetate and glucose. Finally, Pa 215 grown on CSP GLAS preferentially catabolized the top tier amino acids, followed by succinate, lactate, acetate, and ultimately glucose. CSP media formulations also contained small concentrations (3 mM) of citrate, which was added as an ion chelator; however, the citrate was readily catabolized as a preferred substrate. Citrate was oxidized during the first growth phase in all tested media. No or minimal overflow metabolism (<4 mM, ∼3 % of lactate and glucose carbon moles in CSP GL medium) was observed. An exception was CSP GLAS grown cultures which accumulated acetate (∼10 mM) above the initial medium concentrations. Upon exhaustion of succinate and lactate, the acetate was catabolized prior to glucose catabolism. The glucose was not completely catabolized in CSP GLAS medium because the medium was nitrogen limited (supplemental material S1). Culture parameters are summarized in Table 1 and data is available in supplemental material S2-S6.

**Figure 2.**
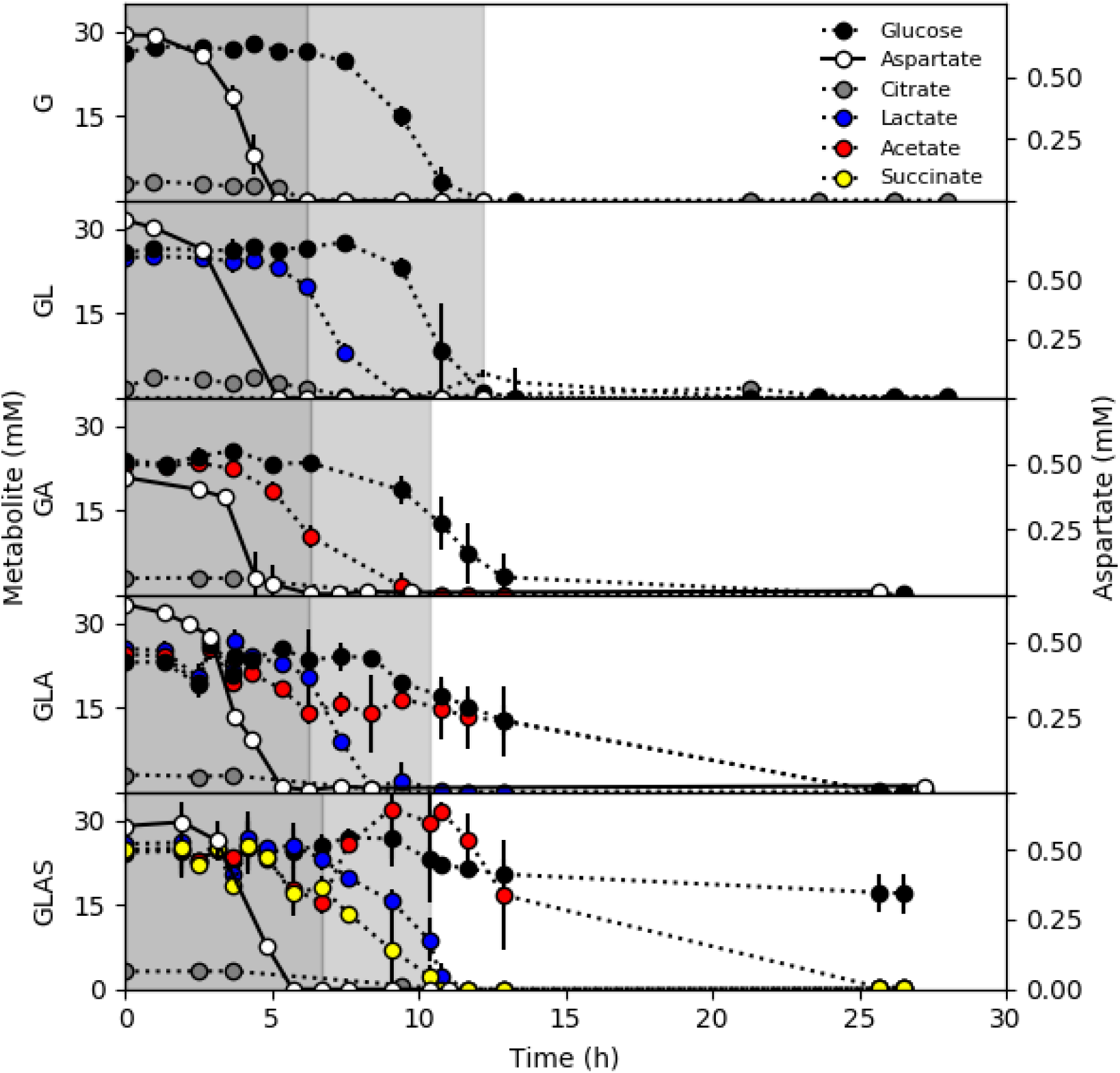
*P. aeruginosa* 215 substrate preference during batch growth in chemically-defined media. Panels plot substrate concentrations as a function of time for five medium configurations supplemented with different carbon sources at a concentration of (22 mM each). G = glucose, GL = glucose and lactate, GA = glucose and acetate, GLA = glucose, lactate, and acetate, GLAS = glucose, lactate, acetate, and succinate. Aspartate concentration (second y-axis) is plotted as a representative top tier amino acid. Cultures demonstrated two exponential growth phases highlighted with different background shading. Medium composition and additional data for each culture can be found in the supplemental data. All values are averages of three biological replicates and two technical replicates, n=6.

The order of amino acid catabolism was assessed for all five CSP medium formulations by binning the amino acids into three categories based on their time of exhaustion (Table 2). All amino acid data including bin designations can be found in the supplemental material S7. The preferred amino acids, referred to here as top tier, included aspartate, asparagine, glutamine, glutamate, and alanine. Most top tier amino acids were binned consistently as quantitated by their small standard deviations. The middle tier amino acids had more variability which may have been CCR-related or based on the temporal granularity of the sampling schedule. The temporal trends in medium pH reflected the metabolism of different substrates. Catabolism of amino acids increased medium pH based on nitrogen chemistry. Catabolism of organic acids also raised the culture pH because the bacterium imports the protonated base, removing a proton from the medium.

Approximately 90% of anabolic nitrogen in CSP G medium was in the form of amino acids (supplemental data S1). *P. aeruginosa* can use ammonium as the sole nitrogen source [38]. CSP G medium formulations were modified with the addition of 2 g/L ammonium chloride to test the effect of nitrogen form. The rCCR phenotype was not changed by the presence of ammonium. The cultures consumed the amino acids as preferred substrates followed by lactate and then glucose (supplemental material S8). The order of amino acid preference remained largely unchanged (supplemental material S9).

The common laboratory strain of *P. aeruginosa*, PAO1, was also grown on CSP GLAS medium. The PAO1 substrate preferences for organic acids and glucose were the same as Pa 215 and the order of amino acid consumption was very similar to Pa 215 (supplemental material S10).

### Proteomics quantifies a constitutive, respiration-centric metabolism

Proteomic data were collected from CSP G and CSP GL grown cultures. Proteomic data are more predictive of cell function than transcriptomic or genomic data alone because they represent an actual allocation of resources into relatively stable, macromolecular pools [39, 40]. Phenotypes were analyzed using label-free proteomics with mass-spectrometry (MS) of whole-cell lysates collected mid-first, exponential growth phase (4 h), early-second, exponential growth phase (7 h), and late-second, exponential growth phase (11 h). The proteomics data were analyzed with focus on central metabolism proteins associated with catabolizing the available substrates and with producing cellular energy.

Enzymes from the tricarboxylic acid (TCA) cycle and associated auxiliary enzymes had largely, constitutive abundances regardless of the medium formulation and the growth phase (Fig. 3). All TCA cycle enzymes except the membrane-associated succinate dehydrogenase were detected and quantified. Additionally, the enzymes oxaloacetate decarboxylase (PA4872) and PEP synthase which process metabolic intermediates for the TCA cycle, were expressed constitutively. The abundance of ATP synthase subunits was also constitutive. Membrane-associated, electron transport chain enzymes were not detected. It was assumed that the enzymes were also constitutively expressed based on the TCA cycle and the ATP synthase protein abundances and the lack of an overflow metabolism.

**Figure 3:**
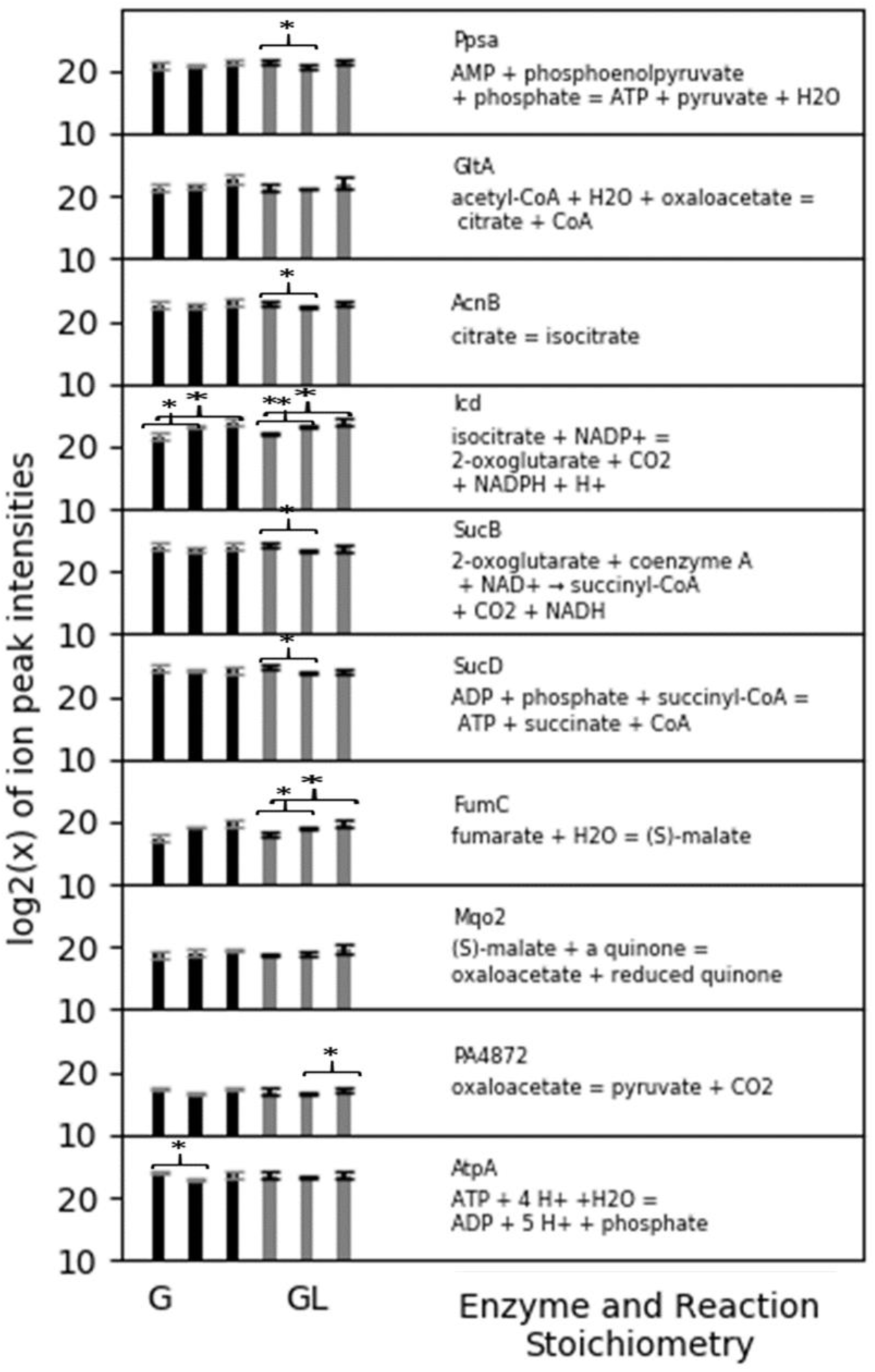
Proteomics data for *P. aeruginosa* 215 grown on chemically-defined, CSP G and CSP GL medium which contained glucose or glucose and lactate, respectively. CSP G culture data are represented by black bars and CSP GL culture data are represented by grey bars. Bars quantify the abundance of the enzyme during the first exponential growth phase (4 h), early second exponential growth phase (7 h), and late second exponential growth phase (11 h). Presented enzymes are involved in the tricarboxylic acid (TCA) cycle, anaplerotic reactions, and ATP synthesis. All values are averaged from three biological replicates. *=p-value <0.05. **=p-value <0.01.

Enzymes associated with the processing of specific substrates did change in abundance based on presence and concentration of substrates, contrary to most TCA cycle enzymes (Fig. 4). Aspartate was plotted as a representative top tier amino acid (Table 2). Protein abundance for aspartate ammonia-lyse (AspA), responsible for the catabolism of aspartate, was elevated during the first exponential growth phase. When aspartate was exhausted, the abundance of AspA dropped as the metabolism shifted to other substrates. Following the depletion of top tier amino acids, the presence or absence of lactate was correlated with increasing or minimal abundances of lactate dehydrogenase protein (Lld), respectively. Pa 215 catabolized glucose, like all Pseudomonads, via the Enter-Doudoroff (ED) pathway [38, 41]. Abundance of ED phosphogluconate dehydratase (Edd) increased while glucose was being metabolized, after the exhaustion of top tier amino acids.

**Figure 4.**
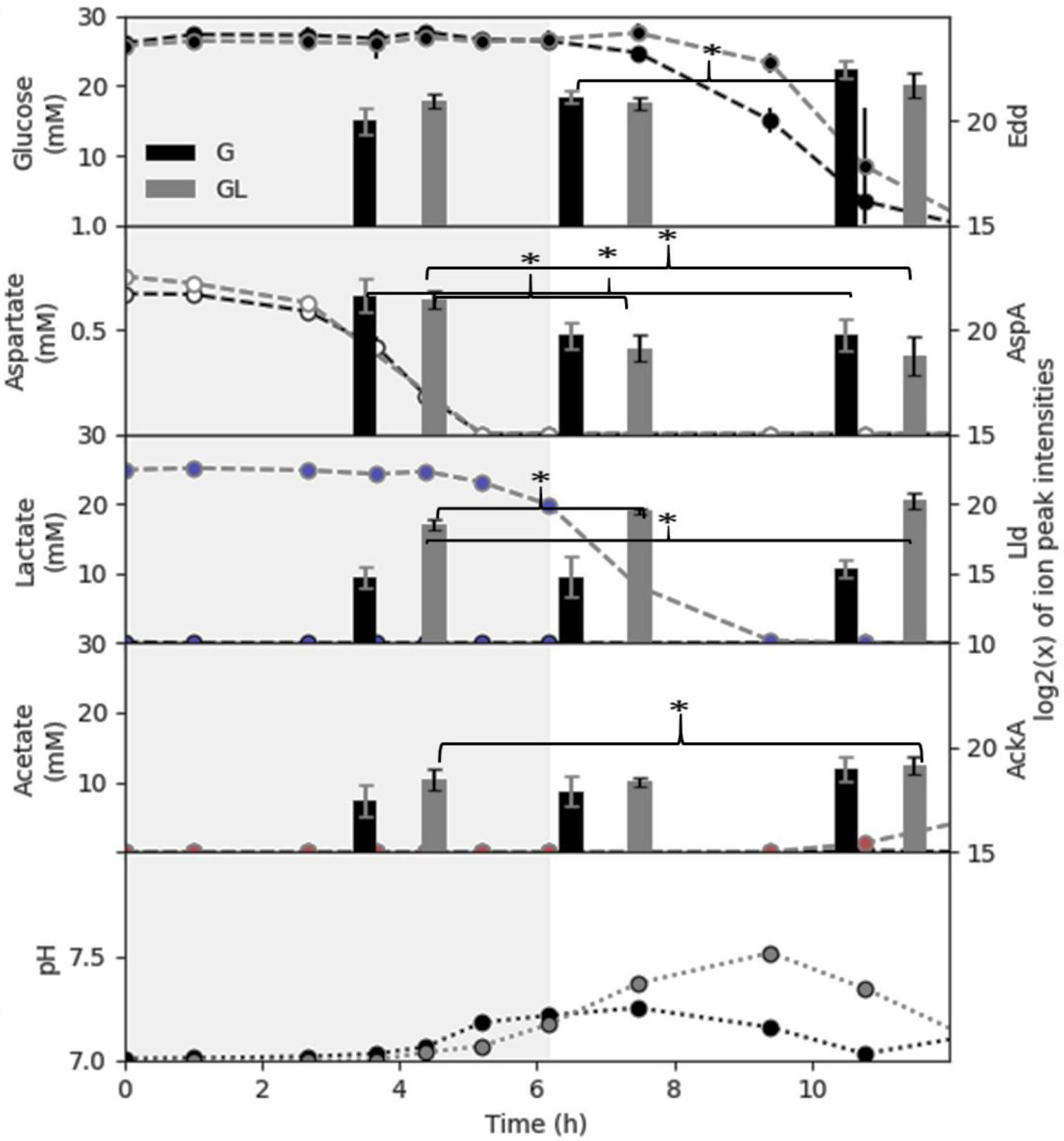
Substrate-specific, protein abundances for *P. aeruginosa* 215 cultures grown on chemically-defined CSP G (black bars and lines) and CSP GL (grey bars and lines) media. Substrate concentrations for each enzyme are plotted in the same panel to highlight relationships. The three bars represent batch growth time points 4, 7, and 11 h. Metabolite values are averaged from three biological replicates and two technical replicates. Protein values are averaged from three biological replicates. *=p-value <0.05. **=p-value <0.01.

Proteomic analysis measured additional proteins that displayed changes in expression during exponential growth and stationary phases. Data can be found at ftp://massive.ucsd.edu/MSV000085590/

### in silico *analysis of rCCR phenotypes*

The order of substrate preference is hypothesized to reflect the ecological strategy used by *P. aeruginosa* to thrive in environmental and medical niches. Computational systems biology tested hypotheses regarding what phenotypic properties were being optimized in the Pa 215 cultures. Analysis utilized a genome-scale, stoichiometric metabolic model of *P. aeruginosa* updated here with genome-supported, amino acid catabolism reactions [42-44](supplemental material S11).

*In silico* testing of ecological strategies was applied first to amino acid preference and included all experimentally measured amino acids except for aromatic and sulfur containing amino acids due to their specialty chemistries. The amino acid preferences did not correlate with the amino acid frequency in genome open reading frames (Fig. 5a) indicating the amino acids were not consumed solely for protein assembly; amino acids were also used as anabolic building blocks for other macromolecules and catabolized for cellular energy. Therefore, simulations considered either the production of cellular energy (*e*.*g*. ATP) or cellular growth which was quantified as carbon moles (Cmol) of biomass.

**Figure 5.**
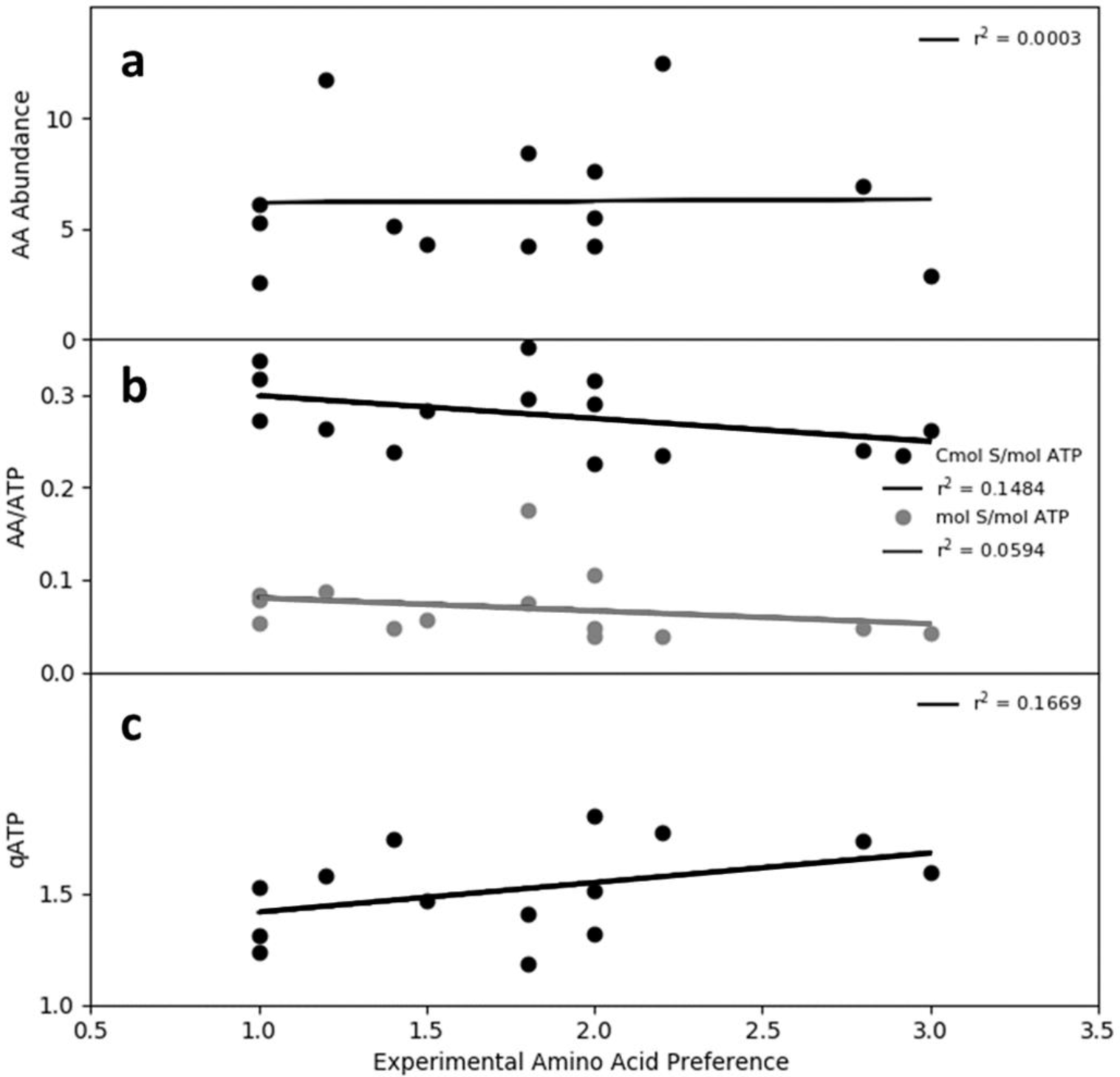
*In silico* analysis of reverse carbon catabolite repression (rCCR) substrate preferences in *P. aeruginosa* 215. a) amino acid frequency in genome open reading frames plotted as a function of the experimental amino acid preference. b) optimal *in silico* cellular energy yield from the catabolism of each amino acid plotted as a function of experimental amino acid preference. Analysis considered both moles of amino acid and carbon moles (Cmol) of amino acid. c) *in silico* maximization of cellular energy rate (qATp = mmol ATP g cdw^-1^ h^-1^) plotted as a function of experimental amino acid preference. Enzyme parameters were based on a survey study of catabolic enzymes (0.5 mmol AA g cdw^-1^ h^-1^, K_m_ = 0.2 mM, and CSP G medium composition, see text for more details)

The first round of *in silico* analyses considered three separate, single dimension optimization criterion 1) maximizing biomass or energy production rates based on electron donor, 2) maximizing product rates based on electron acceptor (O_2_), or 3) minimizing nutrient investment into the proteome required for product synthesis.

### Amino acid preference does not correlate with *in silico* maximization of rates

Computational approaches for studying metabolism often assume cells utilize metabolic potential to maximize growth rate [45-47]. Amino acid preference was analyzed using this theory. Simulations identified the optimal conversion of each individual amino acid into cellular energy or biomass. The *in silico* phenotypes which maximized yields, oxidized completely the substrates, consistent with *in vitro* cultures which did not utilize overflow metabolisms.

Maximizing cellular energy yield for each amino acid, on a mole substrate or Cmol substrate basis, did not correlate with amino acid preference, having r^2^ values of 0.06 and 0.15 respectively (Fig. 5b). The phenotypes which maximized yields were used to calculate maximum product rates using enzyme parameters from survey studies [48-50] and experimental medium composition (supplemental material S1). Maximizing the rate of cellular energy production (or growth) did not predict amino acid preference, as the correlation was r^2^= 0.17 (Fig 5c)(supplemental material S12-S17).

*P. aeruginosa* has a respiration-centric metabolism. The rate maximization criterion was also applied to O_2_ which is required to fully oxidize the amino acids under the experimental conditions. This alternative, rate maximization criterion predicted substantial overflow metabolisms for most of the amino acids. This predicted phenotypic trait was not consistent with the experimental data indicating the criterion was not relevant for Pa 215 metabolism (supplemental material S12).

Amino acids with high cellular energy yields (mol ATP (mol amino acid)^-1^) also had high biomass yields (Cmol biomass (mol amino acid)^-1^); the two yields correlated with an r^2^ value of 0.99 (supplemental data S17). Therefore, maximizing rates for cellular energy production or biomass production had similar trends and neither predicted rCCR phenotype (supplemental material S12).

### Single dimension optimization of resource investment predicts overflow metabolism, inconsistent with rCCR phenotypes

Computational analysis was used to identify phenotypes that minimized resource investment into catabolic pathways. Explicit investment models were not possible in non-model organism, *P. aeruginosa*. Therefore, two previously developed resource proxies were applied to estimate relative, proteome investment into central metabolism [23, 26, 51, 52]. The flux minimization proxy [23, 26, 53] assumes the total network flux is proportional to the enzymatic resources required to synthesize the necessary proteome. Another proxy for protein investment minimizes the number of enzyme catalyzed reactions (a.k.a. minimal proteome investment) identifying the smallest proteome required to realize an *in silico* phenotype [23]. These hypotheses assumed central metabolism enzymes could be approximated as having the same molecular weight with the same amino acid distribution.

Both proxies for resource investment, when applied as the single optimization criterion, predicted overflow metabolisms for most amino acids (supplemental material S14, S15). The *in vitro* experimental cultures did not demonstrate substantial overflow metabolisms indicating this single criterion was not relevant for Pa 215 phenotypes.

### Substrate preference is consistent with a resource utilization strategy optimizing substrate oxidation and proteome investment

Life occurs in multifactorial environments with multiple stressors influencing phenotypes [24, 54, 55]. Two dimensional, optimizations of *in silico* phenotypes were performed where the first dimension considered the optimal conversion of substrate into cellular energy which completely oxidized the substrate. The second dimension quantified the nutrient investment necessary to synthesize the central metabolism proteome (supplemental material S14, S15). Both the flux minimization and minimal proteome investment proxies were tested. Two-dimensional optimization (2-DO) using the flux minimization proxy had poor correlations with the observed amino acid preferences (Fig 6a). Alternatively, 2-DO using complete substrate oxidation and the minimal proteome proxy predicted the experimental preference for amino acids (r^2^ = 0.67) (Fig. 6b).

**Figure 6.**
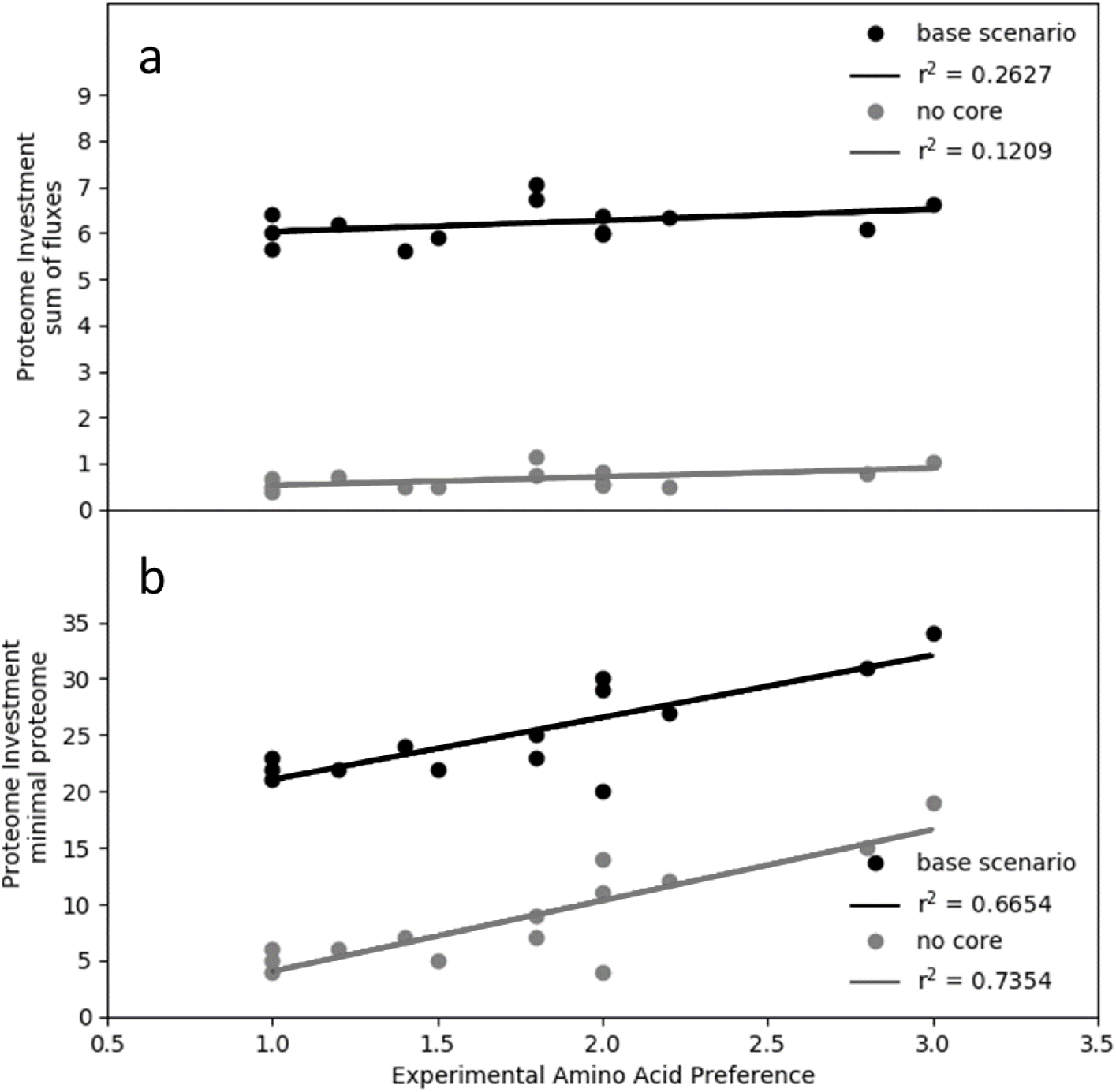
Computational analysis of amino acid preference based on resource investment into proteome for complete oxidation of substrate. a) correlation of sum of fluxes proteome proxy vs. experimental amino acid preference. Analysis considered base metabolism scenario and refined, no core metabolism scenario. See text for more details. b) correlation of minimal proteome vs. experimental amino acid preference. Analysis considered base metabolism scenario and refined, no core metabolism scenario. See text for more details.

2-DO was further refined using experimentally measured proteomics data. The constitutively expressed TCA cycle, anaplerotic enzymes, ATP synthase, and electron transport chain (Fig. 3) were considered part of a core, constitutive proteome, independent of substrate. The refined, 2-DO theory considered only the resource investment in addition to the conserved, core proteome. This theory lead to improved predictions of amino acid preference with the minimal proteome investment theory but not the flux minimization theory (Fig. 6a, 6b). The outlier amino acid in Fig. 6b was serine. Serine is catabolized via the L-serine dehydratase enzyme which is O_2_-labile suggesting higher cell densities and lower O_2_ concentrations are necessary for its functionality [56, 57]. The predictive accuracy of the analysis improved to a correlation of r^2^ = 0.88 if serine data were excluded.

2-DO, considering complete oxidation of substrate and minimal proteome investment, was extended to the other available substrates including organic acids and glucose. Analysis applied the minimal proteome investment with conserved core proteome assumption and considered both cellular energy production as well as the more complex biomass production (Fig. 7a). The *in silico* analysis accurately predicted experimental, substrate preference for amino acids, citrate, succinate, lactate, acetate, and glucose. Correlations had an r^2^ of 0.94 and 0.73 for cellular energy and biomass production, respectively. The biomass simulations included an aggregate amino acid substrate pool, which was not considered for cellular energy simulations, and as anticipated, the aggregate amino acid pool greatly reduced the requirement for enzymatic steps by negating *de novo* amino acid synthesis reactions (Fig. 7a, supplemental material S14, S15).

**Figure 7.**
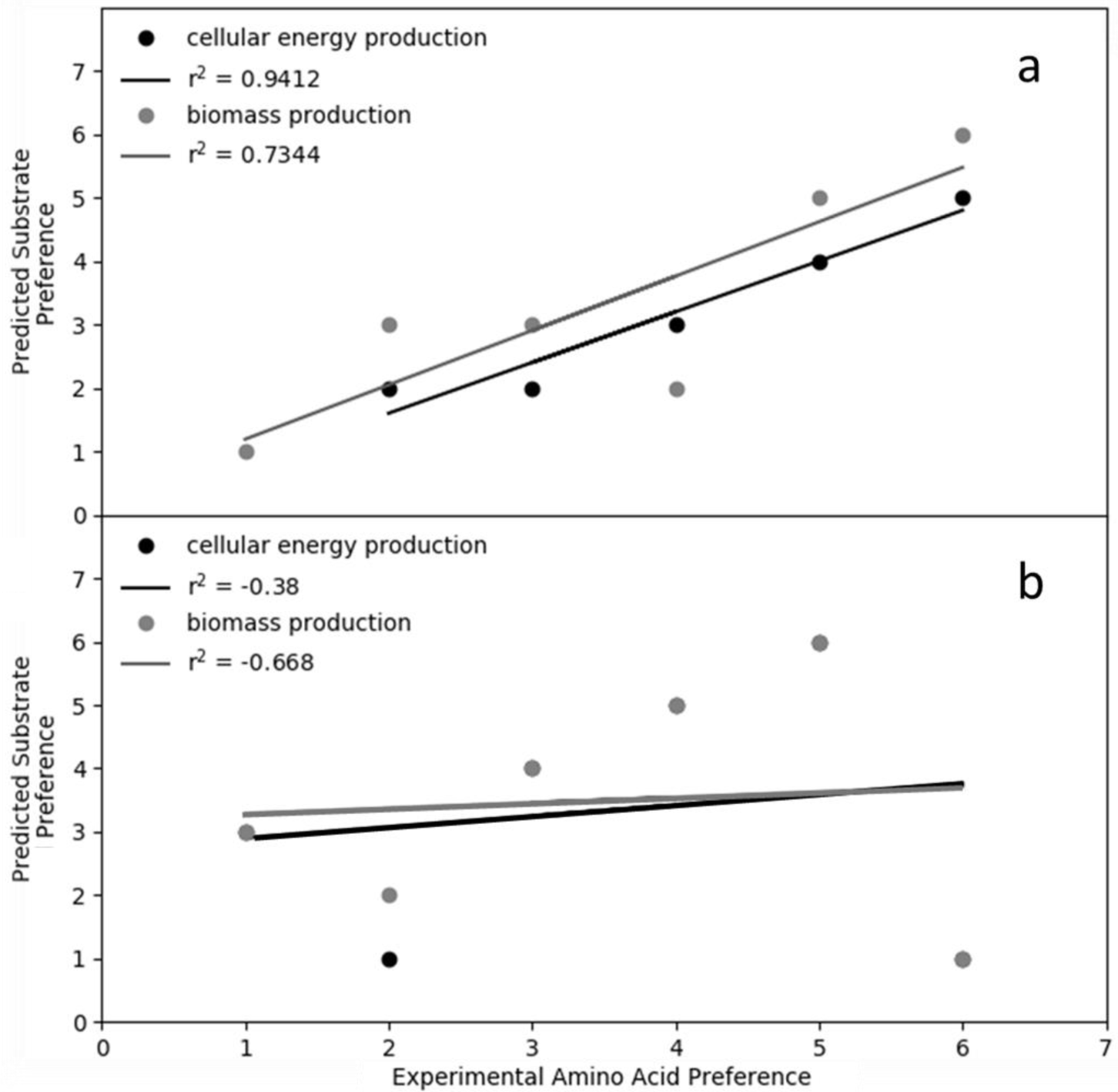
Predicted substrate preference, based on *in silico* analysis, compared to experimental substrate preference for cultures of *P. aeruginosa* 215 growing on chemically-defined media. The experimental substrate preference was: 1. amino acids, 2. citrate, 2. succinate, 3. lactate, 4. acetate, and 5. glucose. a. Two-dimensional, substrate preference predictions for cellular energy production and biomass production. Simulations used the minimal proteome investment proxy and refined, core proteome theory. See text for details. b. Predicted substrate preference based on the maximization of rate criterion for cellular energy production and biomass production. *In silico* product yield on substrate was assumed proportional to rate.

The maximization of rate criterion was also tested with the additional substrates. The analysis assumed optimal product yields on substrate were proportional to the optimal product rates. The maximum rate criterion did not predict the experimental preference for organic acids over glucose. In fact, the predicted preferences had negative correlations with the experimental data (Fig. 7b). Additional optimizations and aggregate substrate simulations were considered (supplemental material S16-S18). None outperformed the presented approach in terms of accuracy and simplicity.

## Discussion

*P. aeruginosa* preferentially consumes nonfermentable, lower energy substrates, such as acetate over glucose. This ecological strategy has enabled its broad, global distribution in the environment and medical niches including chronic, diabetic ulcers. The hierarchy of substrate preferences for *Pa 215* was: amino acids such as aspartate, followed by citrate, succinate, lactate, acetate, and finally glucose. These preferences were also observed with *P. aeruginosa* PAOI grown on CSP GLAS medium (supplemental material S10). *Pa 215* maintained, constitutively, core TCA cycle enzymes and regulated the abundance of the proteins required for specific substrates as needed to convert the substrates into central metabolism intermediates. Analysis using an *in silico* metabolic model determined the rCCR phenotype was consistent with a multidimensional, resource utilization strategy where substrate preference was based on minimizing the proteome investment required to completely oxidize the metabolite. The rCCR phenotypes were not consistent with the commonly applied, systems biology criterion, which maximizes growth rate [45-47].

Pseudomonads are commonly found in consortia [58, 59]. Consortia with populations expressing rCCR and cCCR phenotypes have the metabolic basis for an effective division of labor. rCCR and cCCR microorganisms have complementary substrate preferences whereby the rCCR population consumes organic acids secreted as overflow byproducts from the cCCR population [60, 61]. Both rCCR and cCCR phenotypes can be predicted using resource investment theories, albeit with the rCCR organism expressing a respiration-centric phenotype and the cCCR organism expressing a glycolysis-centric, overflow phenotype. Division of labor can alleviate overlapping substrate preferences between species avoiding competition [62-68]. Additionally, the CCR-based, division of labor could create a positive feedback mechanism by removing inhibitory organic acids and raising local pH ultimately increasing consortia productivity by permitting a more complete depletion of substrates [69]. Natural environments are often limited by anabolic nutrients including nitrogen [22]. Division of labor can also enable higher consortia fluxes for a scarce nitrogen supply based on the nonlinear relationship between enzyme flux and resource investment [21, 70]. This kinetic effect can translate into consortia having a better metabolic return on limiting nutrients, leading to higher biomass accumulation and higher host bioburden [21, 71-75].

Most virulence mechanisms are nutrient acquisition strategies that are also effective in medical niches [9]. CCR regulates a wide range of social behaviors and likely modulates division of labor which would facilitate substrate acquisition by rCCR microorganisms [21]. *P. aeruginosa* preference for non-fermentable substrates like succinate makes it a secondary resource specialist that requires terminal electron acceptors like O_2_ or nitrate [76, 77]. However, O_2_ is often limiting in biofilms where cellular O_2_ consumption rates are faster than diffusion rates [37, 78]. *P. aeruginosa* possesses effective mechanisms to acquire scarce resources like O_2_ [75, 79-83]. For example, *P. aeruginosa* secretes a cocktail of moieties such as pyocyanin, quinolones, and cyanide [58, 84-90]. Exposure to this cocktail can manipulate the *S. aureus* cCCR phenotype, driving it toward overflow and fermentative metabolisms [75, 91, 92]. Collectively, the compounds enable a secondary consumer to influence the metabolism of neighboring cells directing their phenotypes toward secreting preferred substrates including organic acids while reserving the O_2_ for *P. aeruginosa* [74, 75, 90].

Lactate has remarkable connections to *P. aeruginosa* substrate preference and medical niches including diabetic wounds. Elevated lactate levels found in diabetic wounds come from two sources. First, diabetic patients can have higher levels of serum lactate due to diabetic ketoacidosis, and secondly, lactate is associated with wound bed colonization by bacteria which produce it as a byproduct [93-95]. > 80% of chronic wounds are colonized by *P. aeruginosa* [96] while 90% of chronic leg ulcers are colonized by *S. aureus* [80, 97-102] which displays a cCCR phenotype [103, 104]. Not surprisingly, these bacteria are often co-isolated [76, 92, 105].

Wounds colonized by multispecies can be more difficult to treat and can have more negative outcomes than wounds colonized by a single species [74, 98, 101, 106]. Synergistic interactions in consortia based on complementary rCCR and cCCR metabolisms could lead to emergent properties such as enhanced biomass productivity based on enhanced resource acquisition and better metabolic return on investment of scarce nutrients, ultimately leading to greater virulence. Mitigating these consortia, through rational countermeasures, will require quantitative knowledge of the metabolic organization which forms the bases of all virulence mechanisms.

## Materials & Methods

### Bacterial strain and cultivation

All experiments used *P. aeruginosa* str. 215, a clinical isolate obtained from a chronic wound at the Southwest Regional Wound Care Center in Lubbock, Texas, USA [107]. Frozen stocks of *P. aeruginosa* 215 were prepared by growing cultures in 10 mL of 1/10 strength tryptone soy broth (TSB) at 37°C with shaking (150 rpm), adding 3 mL of 20% glycerol, mixing well, and making aliquots of 1 mL volumes then stored until use at −80°C.

Frozen stocks were plated on tryptic soy agar (TSA) at 37°C for 12 h, five colonies were picked to inoculate 10 mL of *Clostridium, Staphylococcus, Pseudomonas* (CSP) medium in culture tubes. CSP is a chemically defined medium developed to support the growth of *P. aeruginosa, Staphylococcus aureus*, and *Clostridium perfringens* as monocultures or consortia [37]. CSP consists of 1.7 g/L Yeast Nitrogen Base without Amino Acids and Ammonium Sulfate (BD Difco(tm)), 0.7 g/L sodium citrate, 0.1 g/L EDTA tetrasodium salt, 100 mL MEM Non-Essential AA Solution (Thermo Fisher Scientific/Life Technologies), 50 mL MEM AA Solution (Thermo Fisher Scientific/Life Technologies), 4.74 g/L KH_2_PO_4_, 8.208 g/L Na_2_HPO_4_, 0.147 g/L glutamine, 2.00 ug/L B_12_, 2.80 mg/L FeSO_4_·7H_2_O, 1.20E-5 g/L CoCl_2_·6H_2_O, and 0.02 g/L of each the following nucleosides: adenine, uracil, cytosine, guanine. For CSP supplemented with one or more organic acids, 22 mM of each organic acid specified was added: CSP G, CSP GL, CSP GA, CSP GLA, and CSP GLAS were supplemented with glucose, glucose and lactate, glucose and acetate, glucose and lactate and acetate, and glucose and lactate and acetate and succinate, respectively.

A total of three culturing tubes containing 10 mL of CSP were each inoculated with about five colonies from the TSA plates, incubated at 37°C with shaking at 150 rpm (tubes were placed at a 45° angle in the shaker to increase mixing) and grown until the cultures reach an OD_600_ of 0.5. 1 mL of each culture was then added to 49 mL of fresh CSP medium in 250 mL baffled flasks giving a culture volume to flask volume ratio of 1:5 and an OD_600_ reading of 0.010. The baffled flasks were capped with gas permeable foam lids and incubated at 37°C with shaking at 150 rpm. Sampling occurred about every hour during the first 12 h and less frequently afterwards.

### Culture sampling

Samples were drawn from each flask for OD_600_, pH, amino acid, and carbon metabolite measurements. An aliquot of 1.5 mL of culture was collected at each sampling, cells were separated from the supernatant using centrifugation at 7000 rpm for 10 min (Eppendorf 5415D microcentrifuge). Supernatants were then filtered using 0.22 μm syringe filters prior to being stored at −20°C.

At each sampling, a volume of culture was collected for OD_600_ measurement. This volume was discarded after measurement. OD_600_ readings were blanked with fresh CSP and samples were diluted, if necessary, to keep OD_600_ measurements ≤ 0.30.

### Organic acid and sugar analyses

HPLC analysis of select carbon metabolites including glucose and organic acids was performed with an Agilent 1200 series HPLC equipped with a variable wavelength detector (VWD) and a refractive index detector (RID) and an Aminex HPX-87H ion exclusion column, 300 mm x 7.8 mm. A mobile phase of 5 mM H_2_SO_4_ was run at a flow rate of 0.6 mL/min for 25 min/injection. A volume of 200 μL of sample was added to an HPLC vial with 200 μL of an internal standard of 1 g/L fucose dissolved in 10 mM H_2_SO_4_. Samples were then loaded into an autosampler, and each was injected twice for a total of two technical replicates for each of the three biological replicates for each time point sampled.

### Amino acid analysis

HPLC analysis of amino acids was performed with an Agilent 1100 series equipped with a diode array detector (DAD) and a ZORBAX Eclipse XDB-C18 column, 4.6 mm ID x 250 mm (5 μm) 80 Å. This setup was used with the Agilent protocol for HPLC analysis of amino acids [108].

### Cell dry weight measurement

A correlation curve between OD_600_ and grams of cell dry weight (g CDW) per liter was constructed. 5 mL aliquots of *P. aeruginosa* culture harvested in mid-exponential growth phase diluted to a range of densities were dried at 80°C for 24 h in aluminum drying pans and weighed. Correlation equation was: (g CDW/L) = (OD_600_) / 2.2256.

### Proteomics analyses

*P. aeruginosa* cultures (n = 3) grown in CSP and LCSP were sampled during exponential growth. Cells were centrifuged at 3600 × g and washed three time with phosphate buffered solution to remove residual media. Cells were resuspended in 1.25 mL radioimmunoprecipitation assay buffer consisting of 12.5 μL of protease inhibitor (Halt Protease Cocktail Inhibitor, Thermo Fisher Scientific, Rockford, IL) to prevent enzymatic degradation upon cell lysis, 0.1 mg/ml lysozyme to solubilize the cell peptidoglycan layer, and 5 mM dithiothreitol (DTT) to cleave protein disulfide bonds. Cells were lysed mechanically in a beadbeater (Mini-beadbeater-1, Biospec Products, Inc, Bartlesville, OK) at 4800 oscillations/min with the remainder of the vial filled with 0.1 mm diameter zirconia/silica beads for a total time of 2.5 min (five cycles at a duration of 30 s each with chilling in an ice water bath between cycles).

Protein concentrations were determined by protein assay kit (DC Bradford Reagent, Thermo Fisher Scientific). 15 μg of proteins were taken from all samples and transferred to centrifugal filter units (Microcon-30kDa Centrifugal Filter Unit with Ultracel-30 membrane, Millipore Sigma, Billerica, MA). The samples were then processed following the filter aided sample preparation method [109], in which proteins were reduced with DTT, alkylated with iodoacetamide to prevent disulfide bond reformation, then enzymatically digested overnight into peptides with trypsin at 1:50 enzyme:substrate (w:w) at 37°C. The resulting peptides were desalted using a C18 column (Macro SpinColumn, Harvard Apparatus, Holliston, MA), dried in a centrifugal evaporator, then resuspended in 5% acetonitrile with 0.1% formic acid to a concentration of 0.2 μg/μL.

200 ng samples of peptides were separated in a high performance liquid chromatography system (1260 Infinity LC System, Agilent, Santa Clara, CA) and a C18 column (3.5 μm particle size, 150 mm length × 75 μm internal diam, Zorbax 300SB, Agilent) using a 60 min mobile phase gradient ranging from 5 - 85% organic (0.1% formic acid in water to 0.1% formic acid in acetonitrile) at 250 nL/min flow rate for a total run time of 75 min. Following LC separation, peptides were ionized by nanoelectrospray with a spray voltage of 1.90 kV and 275°C capillary temperature, then analyzed in a high resolution Orbitrap mass spectrometer (Orbitrap Velos Pro purchased in 2007, Thermo Scientific, Waltham, MA) with automatic gain control set at 10^6^ ions and injection times of 1 - 200 ms. Full scan mass spectra from m/z 400 - 2000 at 30,000 mass resolution were collected in data-dependent acquisition mode. Ten precursor ions were selected from each full mass scan for analysis by tandem mass spectrometry (MS/MS, using 30% energy in HCD mode for fragmentation).

Raw mass spectra data files were processed for protein identification using the MaxQuant software (v. 1.5.3.30) [110] with main search parameters of 4.5 ppm peptide tolerance, 20 ppm MS/MS match tolerance, 10 ppm MS/MS *de novo* tolerance, seven minimum peptide length, carbamidomethyl as fixed modification, 0.01 FDR, oxidation and acetylation variable modification, and enabled search for contaminants. Protein abundances were further data processed and normalized with log transformation (base 2) for data visualization and statistical analyses (ANOVA, p-value <0.05) using the Perseus software (v.1.5.4.0) [111]. The Search Tool for Retrieval of Interacting Genes (STRING) database (v. 10.5) [112] was used for protein-protein interactions amongst the statistically significant proteins, set at medium confidence of 0.4 and protein annotation (functional enrichment analyses). The Kyoto Encyclopedia of Genes and Genome (KEGG) database was also used to visualize lactate and glucose metabolism [113].

### *In silico* analysis of metabolism and resource allocation

A genome-scale, stoichiometric model of *P. aeruginosa* (iMO1086) [42-44] was run with COBRA Toolbox (https://opencobra.github.io/cobratoolbox/stable/cite.html) in MATLAB using the Gurobi optimization program (http://www.gurobi.com) (supplemental material S11, S19). Carbon and O_2_ limitations were modeled by setting the carbon (5 mmol/g/h) or O2 (20 mmol/g/h) uptake rates, respectively, for each of the examined carbon sources and maximizing the production of biomass or cellular energy (i.e. quantified as the number of ATP bonds hydrolyzed). Enzyme limitation was modeled by minimizing the number of participating reactions for specified substrate uptake rates while producing biomass or cellular energy. Suboptimal solutions between the minimal total flux and the maximum product yield (biomass or cellular energy) or the minimum proteome and the maximum product yield (biomass or cellular energy) were identified by minimizing an aggregate objective function. The aggregate objective function was the sum of either total flux or total proteome and a weighted flux through the substrate transport reaction of interest. Optimization between proteome investment and product yield was achieved by changing the weight of the flux through the carbon or oxygen transport reaction of interest. The algorithms can be found in supplemental material S19.

## Acknowledgements

This work was supported by the National Institutes of Health award U01EB019416 and the Army Research Office award W911NF-16-1-0463.

## Author contributions

Conception: LH, MAH, RPC

Design of work: SLM, KAH, LH, RPC

Acquisition and analysis: SLM, YY, KAH, RPC

Interpretation of data: SLM, YY, KAH, MAH, LH, RPC

Drafting and revising of document: SLM, YY, KAH, MAH, LH, RPC

The authors declare no competing interests.

